# Shrinkage improves estimation of microbial associations under different normalization methods

**DOI:** 10.1101/406264

**Authors:** Michelle Badri, Zachary D. Kurtz, Richard Bonneau, Christian L. Müller

## Abstract

Consistent estimation of associations in microbial genomic survey count data is fundamental to microbiome research. Technical limitations, including compositionality, low sample sizes, and technical variability, obstruct standard application of association measures and require data normalization prior to estimating associations. Here, we investigate the interplay between data normalization and microbial association estimation by a comprehensive analysis of statistical consistency. Leveraging the large sample size of the American Gut Project (AGP), we assess the consistency of the two prominent linear association estimators, correlation and proportionality, under different sample scenarios and data normalization schemes, including RNA-seq analysis work flows and log-ratio transformations. We show that shrinkage estimation, a standard technique in high-dimensional statistics, can universally improve the quality of association estimates for microbiome data. We find that large-scale association patterns in the AGP data can be grouped into five normalization-dependent classes. Using microbial association network construction and clustering as examples of exploratory data analysis, we show that variance-stabilizing and log-ratio approaches provide for the most consistent estimation of taxonomic and structural coherence. Taken together, the findings from our reproducible analysis workflow have important implications for microbiome studies in multiple stages of analysis, particularly when only small sample sizes are available.

## INTRODUCTION

Recent advances in microbial amplicon and metagenomic sequencing as well as large-scale data collection efforts provide samples across different microbial habitats that are amenable to quantitative analysis. Following the organization of sequence data into Operational Taxonomic Units (OTUs) or Amplicon Sequence Variants (ASVs), via pipelines such as qiime (1), mothur (2), or dada2 (3), the resulting count data are then available in tabular format for statistical analysis. Downstream analysis tasks include assessing community diversity (4), differential abundance analysis, associating bacterial compositions to system-specific ecological and biomedical covariates, and learning microbe-microbe associations.

However, technical artifacts inherent in microbial abundance data preclude the application of such analysis tasks directly on the measured counts. The data typically comprise a high proportion of zeros and carry only relative information about species abundance. The total number of read counts for any given observation is limited by the total amount of sequencing, quality of DNA preparations, and other technical factors and does not represent the community abundance or total species abundance in the sample or ecosystem. For example, unequal amplicon library sizes can bias sequencing reads to taxa from the larger sample, regardless of true abundance profiles. Although some recent studies have used controlled communities, spike-in controls and other innovations to obtain total community size (5–8), in the majority of experimental designs, the community size is unknown, and, thus, our data is best thought of as containing relative or compositional information (each OTU fraction of total counts, total community size unknown) (9, 10). Additionally, technical variation due to sequencing such as differences in amplification biases and batch effects due to multiple sequencing runs can hamper proper quantification of microbial compositions (11).

To ameliorate these biases, general data normalization methods have been proposed to correct for sampling bias, library size, and technical variability, including workflows from RNA-seq pre-processing and compositional data analysis (12–15). Dedicated normalization and modeling strategies are also available for specific analysis tasks, most prominently, for differential abundance testing (16–19).

Here, we examine data normalization schemes in connection with a fundamental multivariate statistical estimation task: inferring pairwise linear associations from microbial count data. Two common strategies that have been adopted for microbial relative abundance data are Pearson or rank-based correlations after data normalization (20, 21) and proportionality (22, 23) as association measure for compositional data. Consistent correlation or proportionality estimation is of paramount importance for a host of downstream analysis tasks, including state-of-the-art diversity estimation that takes the connectivity of the community into account (4), direct microbial association network inference (12), discriminant analysis, and microbial community clustering.

While previous work (20) has assessed the precision of correlation detection strategies on synthetic microbial sequencing count data, we took a different approach and investigated the behavior of linear association estimation on the largest-to-date citizen-science sample collection, the American Gut Project (AGP) (24). The large available sample size *n*>9000 allows us, for the first time, to critically measure the asymptotic consistency of combinations of data normalization and association estimation techniques. More specifically, given the lack of “gold standard” microbial associations in gut microbial communities, we evaluated different estimation strategies on subsamples of the AGP data of increasing size. We asked the question whether and how association patterns inferred from small but realistic sample sizes of tens to a few hundreds of samples resemble those inferred using the entire data set.

Using a comprehensive set of evaluation criteria and summary statistics, we first show that, independent of any specific data normalization scheme, standard linear association measures are unreliable in the small sample regime. We propose the concept of shrinkage estimation (25) as an effective strategy for consistent association estimation in the small sample regime and quantify the effects of sample size on data normalization and association estimates on several down-stream analysis tasks, including microbial association (or relevance) network inference and clustering. Figure 1 shows the proposed analysis framework used in this study.

**Figure 1.**
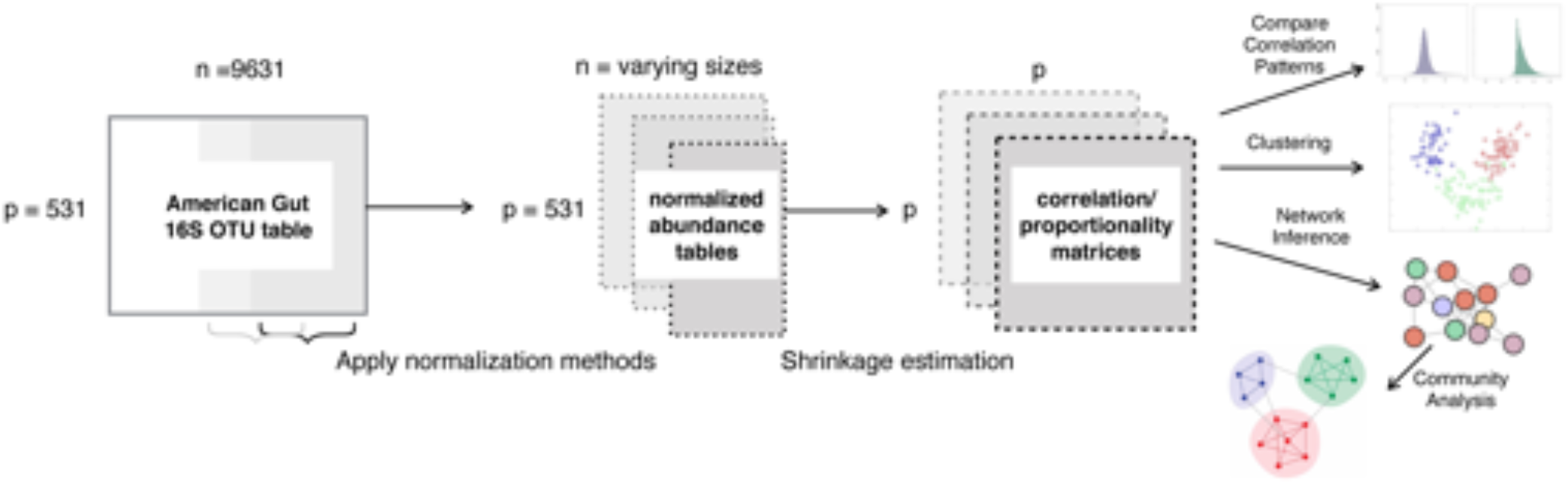
Framework for examining the effects of normalization methods on linear association estimation with increasing sample size. Comparative summary statistics of the resulting association matrices include histogram-based analysis, distance-based matrix comparison, hierarchical clustering, and association network analysis.

Our analysis revealed that all normalization-dependent association estimates in the AGP data can be broadly grouped into five categories and that variance-stabilization and logratio approaches provide the most consistent estimation in terms of taxonomic and community structure coherence. Our findings, available in a fully reproducible statistical analysis R workflow at Synapse ID: syn21654780, have important implications for microbiome studies in multiple stages of analysis, most prominently in the presence of small sample sizes. In particular, we believe that our developed shrinkage estimation framework will improve the consistency of future microbiome data analysis studies at almost no additional computational cost.

## METHODS

To examine the interplay of data normalization and association estimation methods, we first describe the four essential ingredients of our analysis: the processed AGP 16S rRNA data set, the comprehensive list of data normalization methods, statistical estimation of linear associations, and downstream statistical evaluation and analysis tools.

### American Gut Project sample collection

We obtained Operational Taxonomic Units (OTU) count tables and mapping files for unrarefied AGP samples (24) from the project website ftp://ftp.microbio.me/AmericanGut/ag-2017-12-04/. We filtered the dataset to contain only fecal samples whose sequencing depths fall above the 10th percentile and removed taxa that were present in fewer than 30% of all samples. This resulted in a data matrix comprised of *p* = 531 taxa and *n* = 9631 samples.

To investigate the sample size dependence of data normalization and association estimation on this dataset, we generated collections of random subsamples of varying sample sizes, ranging from 25 ≤ *n* ≤ 9000. Large-sample reference association estimates were calculated using a subset of samples at *n* = 9000. To simulate reference data under null correlation or proportionality, we randomly shuffled OTU count data across samples prior to normalization.

### Normalization methods

All normalization methods require as input OTU counts, collected over *n* samples and stored in a matrix 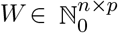. Each row is a *p*-dimensional vector 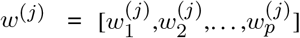, where *j* = 1,…,n is the sample index, 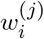 is the read count of OTU *i* in sample *j*, and 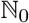 is the set of natural numbers {0,1,2,…}. Let the total OTU count for sample j be 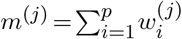. Several methods require the application of a log transformation, thus requiring nonnegative input data. For consistency, we include a pseudocount of 1 to all OTU input data if zero counts are not explicitly handled by the respective normalization scheme. We consider the following data normalization or transformation schemes.

#### Total Sum Scaling

A standard approach for normalizing count data is to divide individual counts by the total OTU counts in a sample, thus scaling the count vector such that the total sum is fixed to 1. This normalization is known as total sum scaling (tss) or total sum normalization. It reads

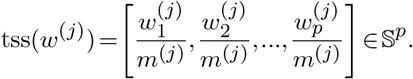

The resulting sample space of the data is thus the (*p*−1)–dimensional simplex.

#### Cumulative Sum Scaling

The tss approach may place unwanted influence on taxa that are highly sampled due to sequencing biases by over-representing it in the scaling factor *m*^(*j*)^ (11). To reduce the influence of these highly abundant taxa for sparse data, cumulative sum scaling (css) has been proposed in (15) and implemented in the metagenomeSeq R package. Rather than normalizing by the total sum, css selects a scaling factor that is a fixed quantile of OTUs counts. Formally,

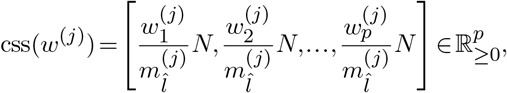

where the scaling factor for sample *j* is 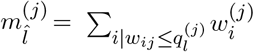. The quantity 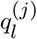 is the sum of read counts up to and including the *l^th^* quantile. *N* is a prespecified constant, e.g., *N* ≜ 1000, chosen such that the resulting data vector resembles the units of the original counts. The sample space of css-transformed data is that of non-negative real numbers 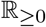.

Let 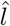 be the index of the 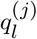 the *l*th quantile for sample *j*, 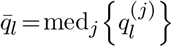 the median 1th quantile across all samples, and let 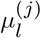 be the mean *1*th quantile. css requires the median absolute deviation of sample quantiles to be empirically stable via the quantity 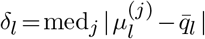. A common choice is to set 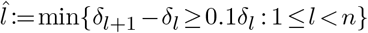 (15). The scaling factor is then defined by summing all the counts up to the smallest value of l that is stable, on average, across all samples that is greater than or equal to the median. The default choice for the median is the lth quantile.

#### Common Sum Scaling

Common sum scaling (com), as introduced in (11), is an alternative to rarefying OTU counts. Counts are scaled to the minimum depth of each sample via

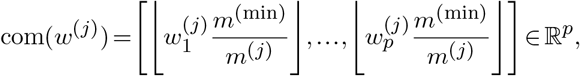

where *m*^(*min*)^ = inf{*m*^(1)^,*m*^(2)^,…,*m*^(*n*)^}. The operator |·| rounds real proportions to the nearest integer.

#### Relative Log Expression

The relative log expression (rle), introduced for gene expression data and available in the DESeq/edgeR package (14). The rle method is defined as follows. Let 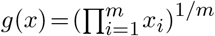 be the geometric mean of an m-dimensional vector *x*, and let 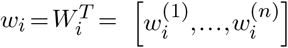 be the vector of counts of OTU *i* over *n* samples (a transposed column vector of count matrix *W*). The numeric scaling factor for sample *j* is

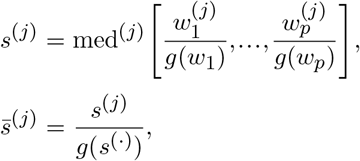

where med[*x*] denotes the median of vector *x* and *s*^(·)^ = [*s*^(1)^,…,*s*^(*n*)^ is a collection of the sample scaling factors. Let the global scaling factor 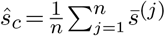 be the arithmetic mean of all normalized scaling factors. The rle is then defined as

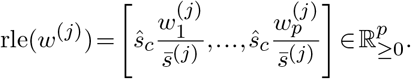

In summary, the rle estimates a median library from the geometric mean over all samples. The median ratio of each sample to the median library is then taken as the scale factor.

#### Inverse Hyperbolic Sine

A standard variance-reducing transformations, often applied to flow and mass cytometry data (26, 27), is the inverse hyperbolic sine function, defined as

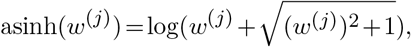

applied element-wise over the sample vector. The resulting data matrix is then mean-centered prior to association estimation.

#### Wrench

The Wrench procedure, introduced in (16), estimates compositional correction factors in the presence of zero-inflation. Wrench is defined as

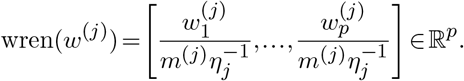

The quantity 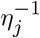 represents a compositional scale factor where 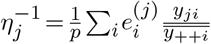. Here, *y_ji_* is the proportion of each feature *i* in sample *j*, and 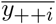 is the average proportion of each feature *i* across all samples. The weight 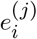 is estimated using the “*W*_2_” scheme, the default choice in the Wrench R package (see also (16) for further details). While Wrench is capable of incorporating information about sample grouping, e.g., for differential abundance testing, we consider all samples to be in a single group.

#### Variance Stabilizing Transform

The goal of variance stabilizing transformations (vst) is to factor out the dependence of the variance on the mean (overdispersion) (14). Consider the mean-dispersion relation 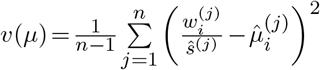. Here, the “size factors” are 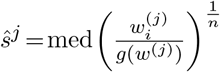 and
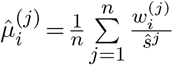 is the sample mean of counts to size factor ratios of sample *j*. The vst is then the integral quantity defined as

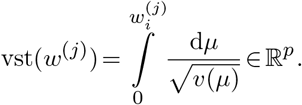

The function *υ*(*μ*) is approximated by a spline function and evaluated for each count value in the column. The vst normalization is available in the DESeq package where the numerical fitting is achieved using local regression on the graph 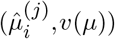. A smooth function *υ*(*μ*) is estimated using an estimate of raw variance: 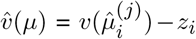 where 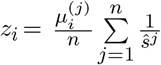 The local regression is parameterized such that large counts are scaled to be asymptotically equal to the logarithm base 2 of normalized counts. When we examined the per-OTU standard deviation (taken across all (*p* = 531) OTUs) plotted against the rank of the average OTU count it can be seen that vst produces similar counts to both clr and a logarithm base 2 transform (Supplementary Fig 1).

#### Centered Log-Ratio transformation

Log-ratio transformations, introduced in (9), transform positive compositional data from the simplex to Euclidean space (9, 12). The centered log-ratio (clr) transform is defined as

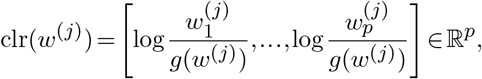

where the ratio is taken with respect to the geometric mean of the composition. The resulting data lies in a *p*−1 hyperplane of *p*-dimensional Euclidean space.

### Estimation of linear associations

Following a transformation of count data under some function 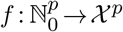 we consider several estimation methods for linear associations among the *p* OTUs.

#### Covariance and Correlation estimation

The standard way of estimating linear associations is the empirical (sample) covariances in the sample space *χ^p^* which forms the basis for many downstream multivariate data analysis techniques, including principal component analysis (PCA), discriminant analysis, metric learning, and network inference.

Formally, column-centering the transformed data results in a *n* × *p* data matrix 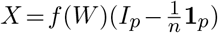 where *I_p_* is the *p*-dimensional identity matrix and 1*_p_* is unit (all-ones) matrix. In matrix notation, the sample covariance matrix (cov) is 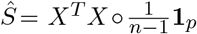, where ◦ indicates element-wise multiplication of two equal size matrices.

The estimate *Ŝ* is a symmetric *p* × *p* matrix with the sample variances along the diagonal and can be normalized to obtain a matrix containing Pearson correlation coefficients. Let 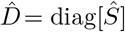 be a diagonal matrix with the p post-transformed OTU variances on the diagonal and zero elsewhere. The Pearson correlation matrix is then 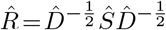. The matrix 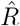 is a symmetric *p* × *p* matrix where each entry 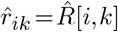 corresponds to the Pearson correlation between OTU *i* and OTU *k* under the data transformation.

The magnitude and sign of the values in 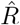 are often interpreted as the association strength and direction, respectively. The sample correlation/ covariance matrices are, however, inadmissible in the *p*≫*n* setting, i.e., when fewer samples than OTUs are available. For example, type I errors may be grossly inflated, since the parameters under estimation are underdetermined. Standard operations for solving systems of linear equations such as PCA are then ill-posed.

#### Shrinkage estimation

To alleviate this shortcoming of the standard sample covariance estimator, regularized covariance and correlation estimation have been developed highdimensional statistics. By imposing structural assumptions about the underlying population covariance, estimators with stronger statistical properties can be derived in the *p*>*n* setting. One prominent structural assumption is sparsity, i.e., only a few strong pairwise correlations are assumed to be present in the dataset. An effective data-driven approach to realizing structural sparsity is shrinkage estimation. While several estimators have been proposed in the literature (28–30), we focus here on Schafer-Strimmer shrinkage estimation (31), as implemented in the R package corpcor. The principle idea of shrinkage estimation is to shrink small sample correlation toward zero where the shrinkage intensities can be efficiently estimated from data (28). More specifically, we estimate individual entries 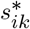 of the shrinkage covariance *S*\* and 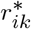 in the shrinkage correlation *R*\* as follows. For all off-diagonal elements in *S*\*, we compute

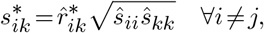

where the shrunk correlation estimates are 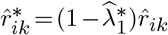. The variances (var) 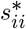 are shrunk in a separate procedure toward the median 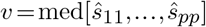 via 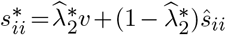.

The shrinkage intensities 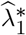 and 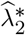 are determined empirically by estimating the variance within the sample covariance matrix (see Supplementary Methods).

#### Proportionality estimation

Covariance and correlation estimation on compositional data has long been criticized due to the necessary presence of negative bias, scale dependence, and subcompositional incoherence in the estimates (9, 32). Association measures based on the concept of proportionality have thus been put forward as an alternative to correlation (22). Here, we consider the symmetric proportionality *ρ_p_* (23, 33) which by default operates on clr transformed data *X^clr^* ≜(*W*). The measure is defined as:

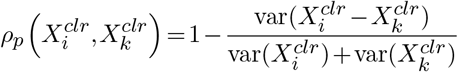

where 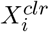 and 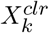 are the columns of the matrix corresponding to OTU *i* and *k*, respectively. *ρ_p_* is a proportionality measure because differences of clr-transformed components are equivalent to log-ratios of compositions. As shown in (23, 33), the measure can be equivalently expressed in terms of covariances and variances as follows:

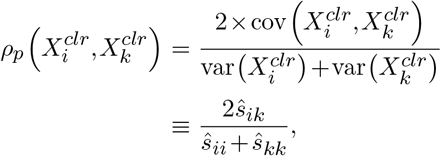

where *ŝ_ik_* are elements of the covariance estimates 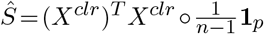 on clr-transformed data. This formulation also clarifies the link between the standard correlation matrix on clr-transformed data and *ρ_p_*: the former uses the geometric mean of *ŝ_ii_* and *ŝ_kk_* in the denominator whereas the latter uses the arithmetic mean. Since *ρ_p_* is completely determined by sample covariances and variances, we expect the measure to have the same drawbacks in the small sample setting as the sample covariance estimators.

We thus propose the novel shrinkage-based proportionality estimator *ρ*_*_ (rhoshrink) as:

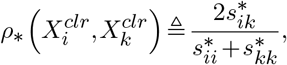

where 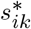 are elements of the Schäfer-Strimmer shrinkage covariance estimator *S*\*.

### Comparing Association Patterns

To quantify similarities of the estimated association patterns across different data normalization method, association measures, and sample sizes, we considered three different distance measures. These distances are then used for comparative low-dimensional embeddings of the different estimates as well as for measuring convergence of the estimators with sample size.

#### Frobenius Distance

Given a pair of *p* × *p*-dimensional association matrices 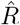 and 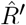, the Frobenius distance measures the sum of squared differences between the corresponding entries and is defined as

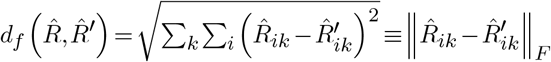

#### Spectral Distance

Given a square, symmetric matrix *A*, let *A* = *UΣU^T^* be its singular value decomposition where Σ is a diagonal matrix with singular values along the diagonal entries, i.e., *Σ_ii_* = *σ_i_*. Let *σ*_max_(*A*) be the largest singular value of *A*. The spectral distance is

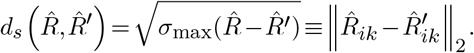

The spectral distance is invariant to unitary transformations of the association matrices.

#### Correlation Matrix Distance

The Correlation Matrix Distance (CMD) (34) measures the orthogonality of two correlation matrices and is defined as

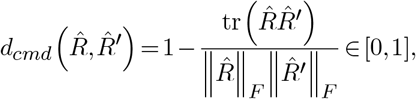

where the trace operator tr(*A*)=Σ_*i*_*A_ii_* is the sum of the diagonal entries of a square matrix.

### Downstream analysis

We considered two downstream exploratory data analysis tasks that require the estimation of microbial associations as input: (i) taxa clustering and (ii) microbial association network construction and community analysis.

#### Clustering

Unsupervised clustering of microbial taxa can help identify microbial sub-communities that may jointly affect host phenotype or reveal experimental and batch effects (24). We considered two popular clustering techniques: spectral and hierarchical clustering.

Spectral clustering requires the construction of an “affinity matrix” from estimated associations. Here, we transformed associations into dissimilarity scores 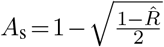 and constructed a *k*-nearest neighbor graph (*k*=2) to obtain a sparse and symmetric affinity matrix *A*. Identification of taxa clusters is based on *k*-means clustering of the first *m* components of the eigendecomposition of of the normalized Laplacian 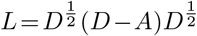, where *D* is the diagonal degree matrix with entries containing the row or column sums of *A* (35). We chose the target cluster size to be number of connected components of the associated affinity graph (35). To assess the taxonomic consistency of a particular clustering, we evaluated the homogeneity of each cluster with respect to the taxonomic families of the underlying OTUs. As a quantitative measure, we computed the ratio of the effective family number (exponential of the Shannon entropy of family counts) to the total number of families detected per cluster.

For hierarchical clustering, we converted association matrices to dissimilarity measures using 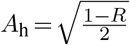 Clustering was then performed using Ward’s method from the hclust package in R. Circular dendrograms were cut using the cuttree method where *k* = 10 was chosen to represent the number of class annotations.

#### Relevance networks and community analysis

Relevance networks (36) are a popular way of visualizing and analyzing the overall structure of the microbial ecosystem. Relevance network construction ranks all pairwise correlation or proportionality values between OTUs by absolute value, selects a certain percentage of highest ranked pairs, and visualizes the resulting set of pairs as edges between OTUs in an association network.

Multiple studies have found a higher prevalence of positive associations between taxonomically related taxa in human gut datasets (12, 37–40). We thus use taxonomic coherence, measured by assortativity (41), as independent summary statistic for relevance networks. When categorical variables are available for each node, the assortativity coefficient of a network takes values in [-1,1] and measures the tendency of adjacent nodes to belong to the same category. In the context of microbial networks, the nodes are OTUs and the associated categories are their inferred taxonomic rank at the Genus level.

We also examined the presence of community structure in the relevance networks using the concept of modularity (42). Similar to clustering analysis, modularity analysis of a network enables the partitioning of nodes into tightly connected sub-communities. Modularity was computed using the fast-greedy algorithm, described in (43) and implemented in the igraph package in R (44).

## RESULTS

Our comprehensive computation and analysis workflow produced several key results which are summarized below. We highlighted statistical properties of association estimation, followed by a comparison of downstream analysis results.

### Shrinkage universally improves consistency of association estimation

We first analyzed the influence of shrinkage on the estimation of associations under different data normalization and sample sizes. We show the convergence properties of association estimation, as measured by Frobenius distance *d_f_* with respect to the large sample limit, with increasing sample size in Figure 2a. Shrinkage universally improves estimates in the low sample regime compared to its sample estimation counterparts. Even when the sample size *n* exceeds the number of OTUs *p*, most shrinkage estimates remain more consistent with their respective large-sample. This behavior is also reflected in the distribution of association estimates at low (*n* = 50) and large (*n* = 9000) sample sizes, as highlighted in Figure 3 for the proportionality measures *ρ_p_* (rhoprop) and *ρ*_*_ (rhoshrink). In the small sample limit, rhoprop produces too extreme estimates compared to the large sample limit (third row in Figure 3). The distributions of rhoshrink estimates in the small and large sample limit were considerably more consistent. This phenomenon was observed for all combinations of data normalization and association estimation (Supplementary Figure 4a, Supplementary Figure 5a). As expected, the influence of shrinkage vanished in the large sample limit, as reflected in decreasing shrinkage intensities with increasing sample size (Supplementary Figure 2).

**Figure 2.**
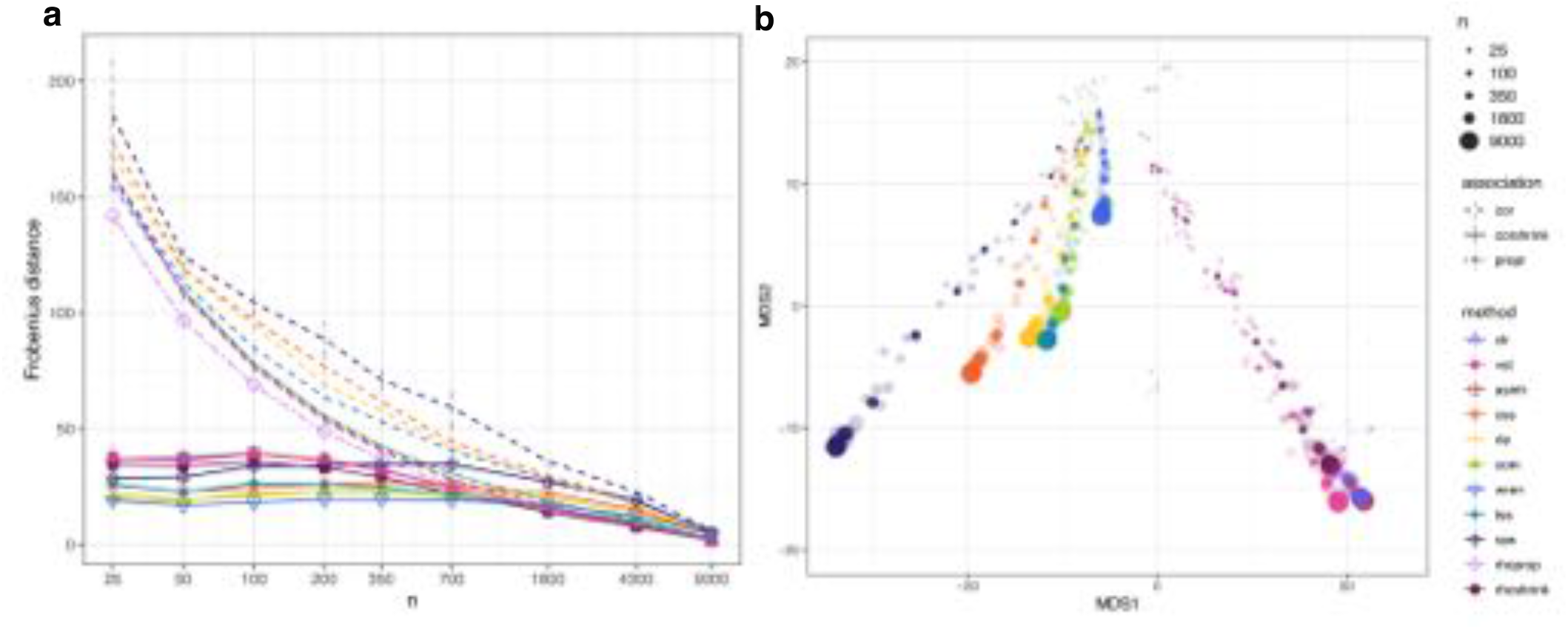
Frobenius distance between estimates of association. a) Frobenius distance between sub-samples of different sizes. Dashed lines represent the mean distance between normalized matrices after Pearson correlation. The solid lines represent the mean distance between normalized matrices where correlation/proportionality estimation with shrinkage was performed. The dot-dash line represents RHO, a proportionality metric. Vertical lines represent standard deviation from the mean for each corresponding method. b) Multidimensional scaling representation of Frobenius distance between correlation structures of varying size estimated from different normalization methods. The most opaque points represent the mean of 5 subsamples of the same size. (color scheme as in A)

**Figure 3.**
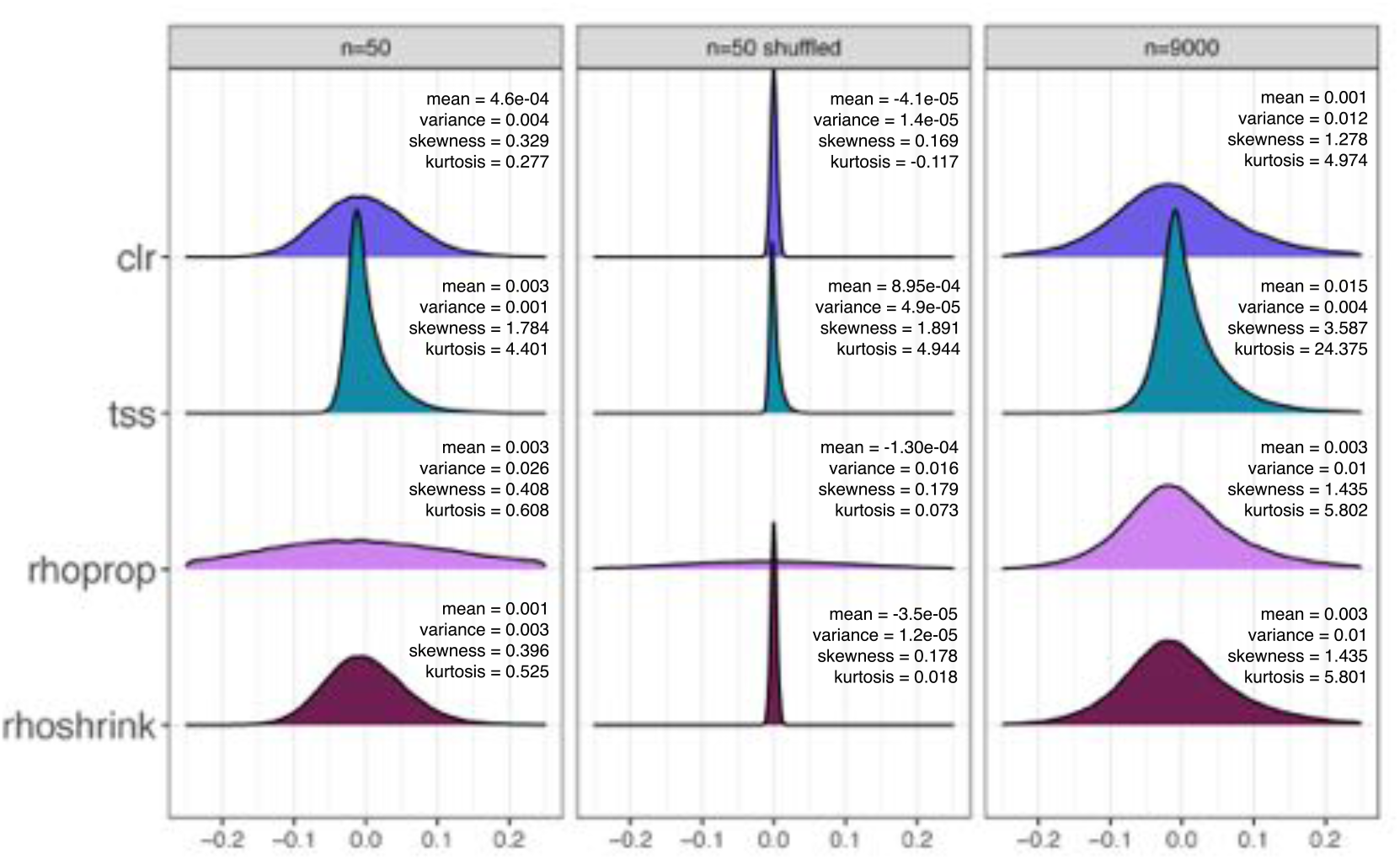
Frequencies of association after transformation and shrinkage. Density plots of association value frequency of matrices after application of different normalization methods at different sample sizes. Mean, variance, skewness and kurtosis are shown for each distribution.

### Normalization methods induce distinct association patterns

We next analyzed the similarity among the different association estimates with increasing sample size using multi-dimensional scaling (MDS). Figure 2b shows a 2D MDS embedding of all shrinkage association estimates using the Frobenius distance. We identified five distinct classes. Association estimates following a variance-reducing/stabilizing transformations (clr, vst, asinh, rhoprop, rhoshrink) form a distinct linear trace in the embedding, ordered along sample size. Correlation estimates on raw count data form another distinct group. Correlation estimates after css and wren normalization form two distinct traces in the embedding. Finally, correlation estimates following the com, rle, and tss normalization form the fifth class of association patterns. For small sample sizes, association patterns are similar independent of the normalization methods. As each of the five classes form a distinct linear trace in the embedding, we used the distances between estimates of different sample sizes to evaluate the rate at which normalization methods arrived at stable patterns of association. In agreement with Figure 2a, we observed that wren, vst, and com arrived at consistent association estimates with the fewest samples, followed closely by rle and tss normalization methods (Figure 2b, Supplementary Figure 3). The observed grouping pattern and convergence behavior is largely invariant to the distance measure used (Supplementary Figure 4b and Supplementary Figure 5b for spectral distance and CMD, respectively).

### Association estimates are positively skewed

We next analyzed the shapes of empirical distributions of shrinkage association estimates for all normalization schemes in three different sample scenarios: Small sample regime (*n* = 50), randomly shuffled data in the small sample regime (*n* = 50), and large sample regime (*n* = 9000). Figure 3 shows clr, tss, rhoprop and rhoshrink distributions across these scenarios (see Supplementary Figure 3 for all others). We observed that variance-reducing/stabilizing transformations (clr, vst, asinh, rhoprop, rhoshrink) induce association distribution that are close to normal with moderate positive skewness. Estimates without shrinkage are considerably wider in the low sample regime (as exemplified for standard proportionality rhoprop vs. rhoshrink in Figure 3). All other normalization schemes induce asymmetric correlation distributions with large positive skewness, resembling correlation distributions on raw count data (Supplementary Figure 3). Positive skewness also persists for association estimates on shuffled data. Although the shapes of shrinkage association distributions are visually similar in the small and large sample limit, we universally observed an increase in skewness and kurtosis with larger sample sizes independent of the normalization scheme.

### Clustering methods are sensitive to normalization and shrinkage estimation

We next focused on analyzing the influence of normalization and association estimation on downstream data analysis tasks. We first considered clustering of OTUs using a large sample limit of *n* = 9000 samples from the AGP dataset. For spectral clustering, we asked the question whether and how normalization and shrinkage influences (i) the standard selection of the number of cluster and (ii) the taxa composition of the resulting clusters. One common strategy for model selection in spectral clustering is the “spectral gap” criterion. The number of selected clusters is considerably larger (*k* ≥ 11) for the variance-reducing/stabilizing transformations (clr, vst, asinh, rhoprop, rhoshrink) than for other normalization methods (*k* ≤ 8)(Supplementary Figure 6). Despite the large sample size, the spectral gap of rhoprop - and rhoshrink-based spectral clustering resulted is still different, resulting in *k*= 11 and *k* = 12 clusters, respectively. The different number of clusters also contributed to marked differences in terms of the homogeneity of taxa compositions, as shown in Figure 4. Variance-reducing/stabilizing transformations produced taxonomically more homogeneous groups at the Family level. rhoshrink-based clustering produced the highest mean cluster purity, indicating strong agreement between estimated OTU associations and taxonomic identity (as shown for Family level in Figure 4 and Supplementary Figure 5). rhoprop- and rhoshrink-based clustering formed very similar but not identical clusters in terms of composition and cluster purity. A larger number of taxa of Family *Ruminococcaceae* and Class *Bacteriodia* cluster together in clr-based clustering compared to tss-based clustering. Taxa clusters derived from css, rle, and wren normalization resulted in no distinct taxonomic grouping (see Supplementary Figure 5).

**Figure 4.**
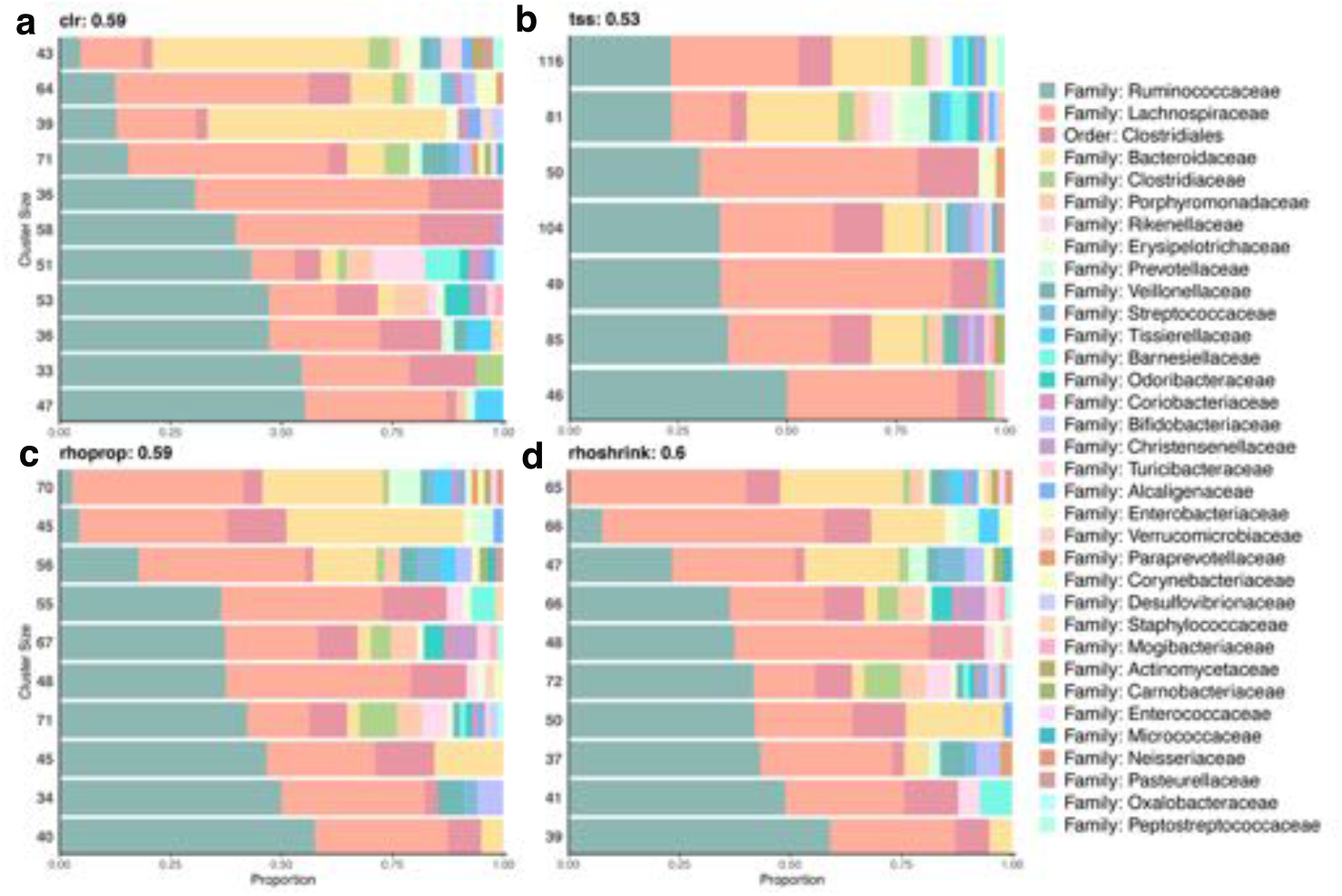
Horizontal stacked bar plot of OTU groups resulting from spectral clustering. a-d) The stacked bar plots represent the composition of OTUs in each cluster at the Family level. Clusters are vertically ordered from highest percentage of the most abundant Family: *Ruminococcaceae*. Horizontally the order represents the highest percentage of Family in each cluster. Numbers on the left axis represent the number of OTUs in each cluster. Values stated next to method name represent cluster purity.

Hierarchical clustering largely confirmed the previous observations. For ease of comparison, we set the number of clusters to *k* = 10 for inference workflows. Figure 5 shows the dendrograms for clr-, tss-, rhoprop-, and rhoshrink-based clustering. While some distinct and homogeneous clusters can be found in the tss case, the majority of taxa has been grouped into a single cluster comprising many families and classes of taxonomically unrelated bacteria. However, taxonomic grouping is well represented by hierarchical clustering of rho- and rhoshrink-based estimates (Figure 5). Similarly, vst and asinh have recovered large groups of the most prevalent family annotation: *Ruminococcaceae*, *Lachnospiraceae*, and *Bacteroidaceae* (see Supplementary Figure 8).

**Figure 5.**
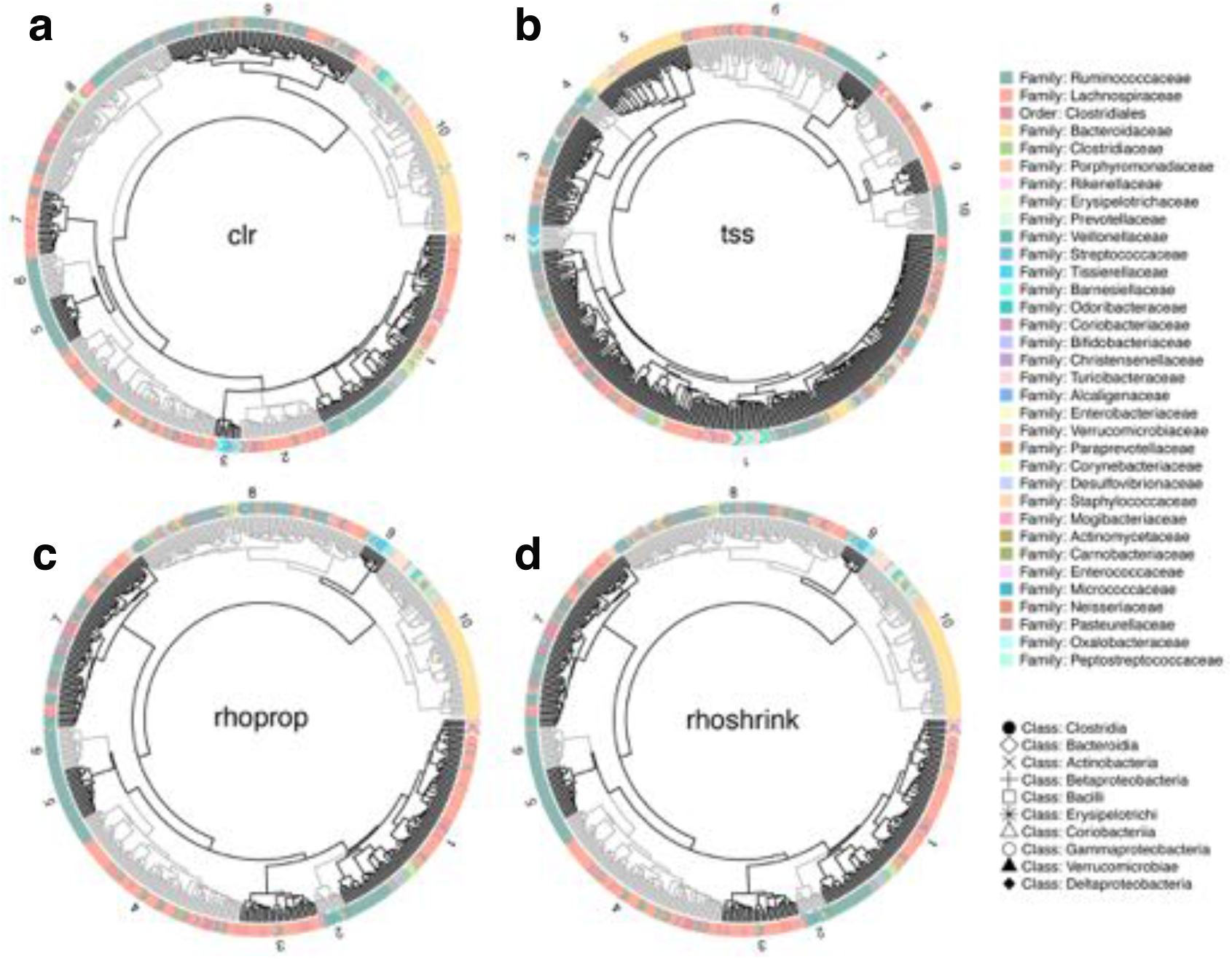
Circular dendrograms of hierarchical clustering patterns amongst taxa Family and Class. Each point surrounding the circular dendrogram represents one of the 531 OTUs in our data set. The color represents Family annotation and shape represents Class annotation. Each dendrogram has been cut hierarchically into 10 trees (representing the 10 Orders to which each Family maps). The grey and black shading is used to highlight different clusters which are numbered.

### Normalization induces relevance networks with different community structures

We next considered the downstream statistical task of learning microbial relevance networks from AGP data. We estimated associations in the large sample limit *n* = 9000 and selected the top 2000 associations for network construction in every data normalization/association estimation workflow. Figure 6 shows network visualizations for clr-, tss-, rhoprop-, and rhoshrink-based relevance networks (Supplementary Figure 9 for the other instances). We identified sub-communities of highly connected taxa using modularity maximization. The number of identified modules ranged between 20 (using Wrench) and 38 (using vst normalization). Relevance networks derived from variance-reducing/stabilizing transformations (clr, vst, asinh, rhoprop, and rhoshrink) were partitioned into 35-38 modules and achieved a maximum modularity score of ≈0.8 (compared to modularity scores of < 0.6 for all other networks). Visual inspection of these networks revealed that members of the *Bacteroidetes* phylum (represented by square nodes in Figure 6) formed tightly connected modules with few edges connecting to other phyla. *Firmicutes* (represented by circular nodes) in networks were divided into a higher number of modules comprising distinct families, including *Lachnospiraceae* (represented by orange circles) and *Ruminococcaceae* (teal circles, Figure 6, Supplementary Figure 9). This striking modularity is less pronounced in the tss-based relevance networks (Figure 6b).

**Figure 6.**
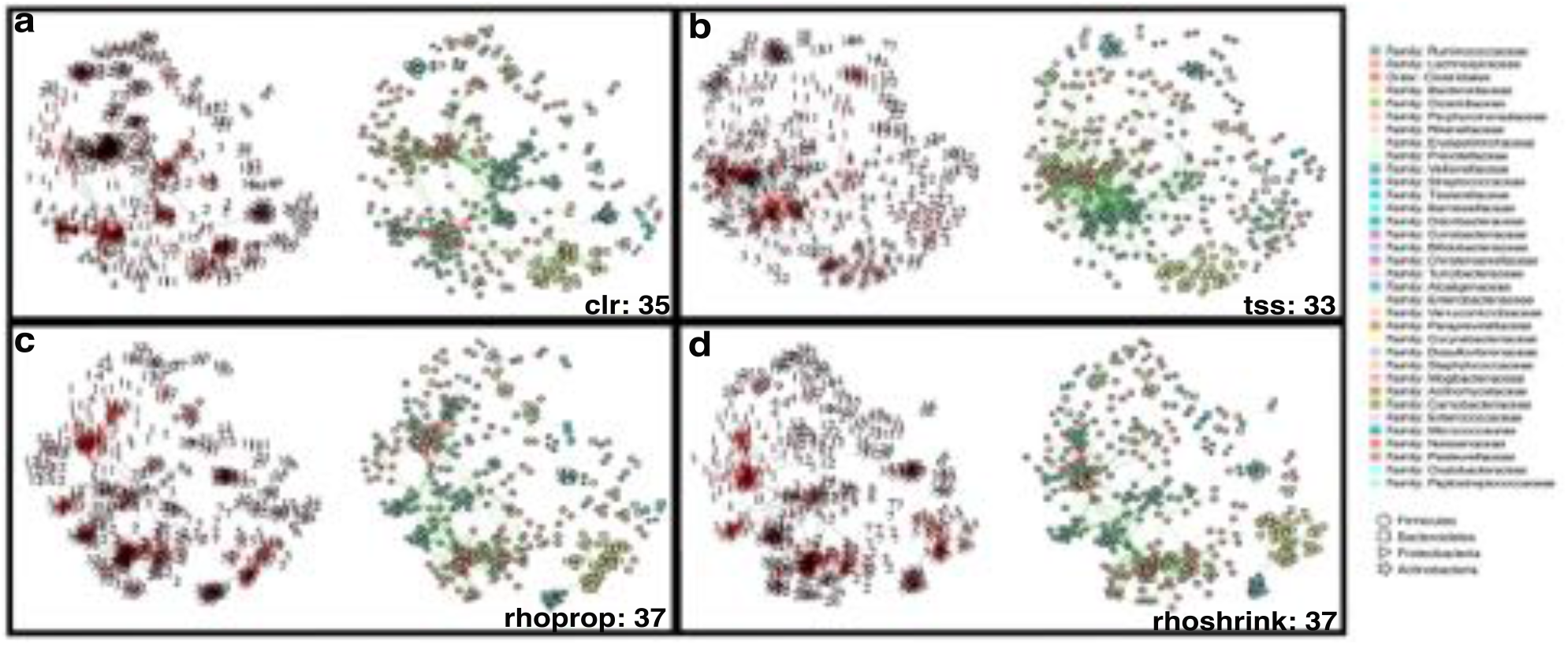
Relevance network visualization displaying modularity. a-d) For networks on the left of each panel every node represents an OTU labeled with module annotation as predicted by the Fast-Greedy modularity algorithm. The networks on the right represent the corresponding taxonomic annotation of the OTU at the family level. Values stated next to method name represent the number of modules in the network. Layout using the force-directed Fruchterman-Reingold algorithm was conserved for both networks in each panel.

Similar to the clustering analysis, we next evaluated the taxonomic coherence of the different networks. Using assortativity on the genus level as a quantitative measure, we found that relevance networks derived from variance-reducing/stabilizing transformations showed the highest overall assortativity in the large sample limit (≈0.35).

We next asked the question whether high-level network properties such as assortativity and modularity were consistent independent of the sample size used to estimate the association networks. We thus repeated the previous analysis for different sample sizes, ranging from *n*= 25 to *n* = 9000. Figure 7 shows the estimated network assortativity and maximum modularity score estimates vs. sample size. We found that for relevance networks derived from variance-reducing/stabilizing transformations, both assortativity and modularity monotonically increase with sample size. Both estimates stabilize around samples sizes *n*≈*p*. For the remaining relevance networks, assortativity estimates monotonically increase with sample size while modularity tends to decrease with sample size. In summary, this analysis implies that estimates of high-level network summary statistics such as assortativity and modularity are inconsistent compared to their large-sample limit.

**Figure 7.**
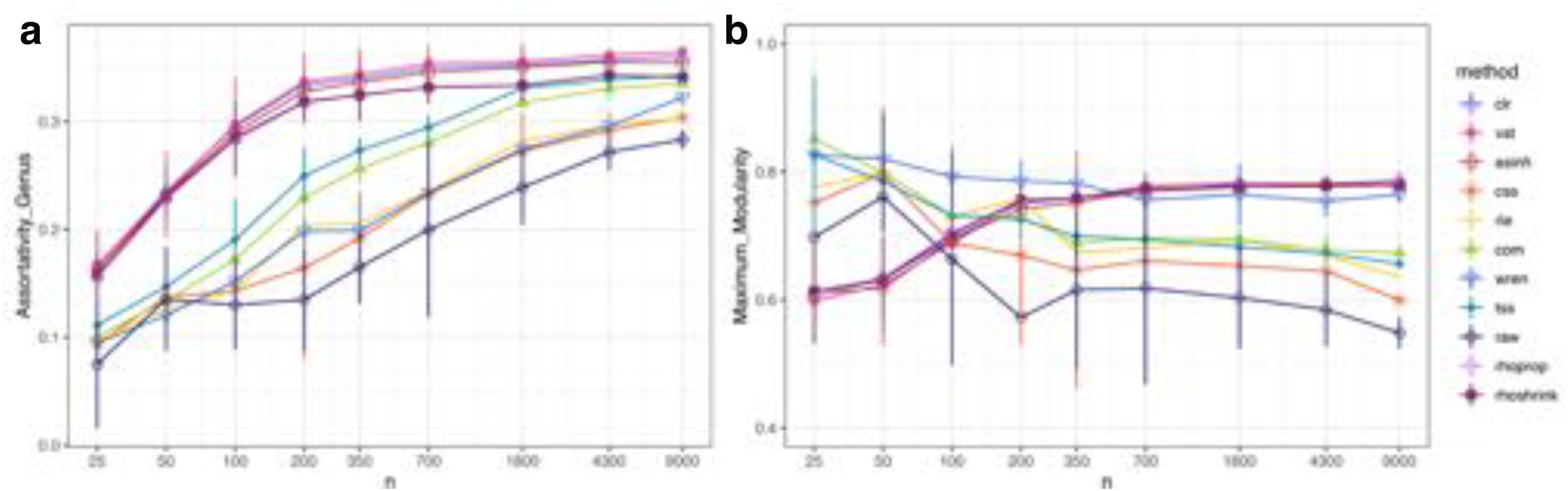
Community analysis of relevance network structure with increasing sample size. a) Assortativity coefficient across sample size of genus annotation. b) Maximum modularity score across sample size at 2000 edges. For all plots lines represent mean and grey ribbon represent standard deviation from the mean.

We next examined the edge overlap from correlation-based relevance networks (clr and tss transformations) and proportionality metrics (rhoprop and rhoshrink). We found a common core of 1086 edges between 349 OTUs that were present in all relevance networks. This consensus network also contained several tightly connected network modules with highly assortative inter-family associations (Figure 8a). Overall, we found that clr-, rhoprop, and rhoshrink-based networks shared the majority of common edges with rhoprop- and rhoshrink-based edge set differing only by a single edge. The tss-based relevance network comprised 779 unique edges not shared by any of the other networks (Figure 8b).

**Figure 8.**
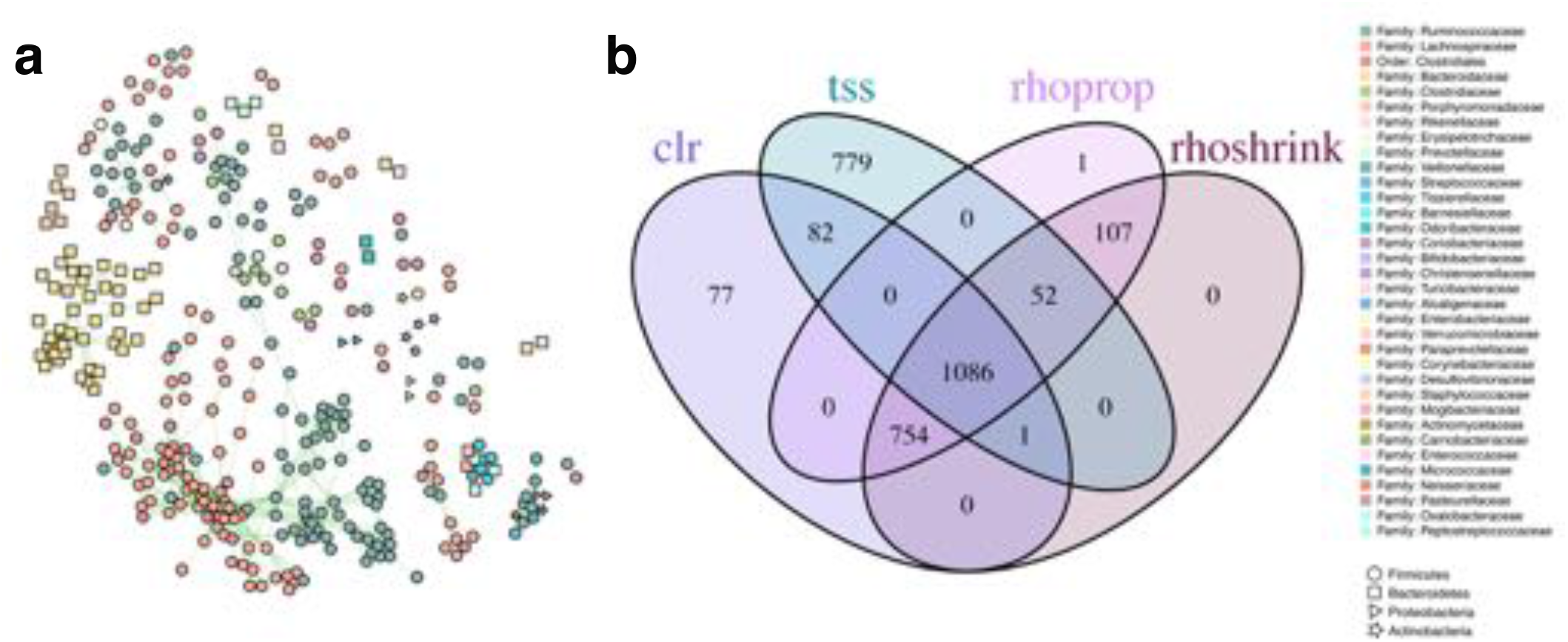
A) Edges in common between relevance networks. This consensus network contains 1086 edges between 346 taxa. Node color represents Family annotation and node shape represents phylum. Network layout was generated using the force-directed Fruchterman-Reingold algorithm. B) Venn Diagram of edges in common from the top 2000 edges between four highlighted normalization methods.

Additionally, we found that clr-, vst-, and asinh-based correlation networks also shared a large common consensus core (Supplement Figure 10a). Similarly, com-, rle-, and tss-based networks showed a large edge set overlap (Supplement Figure 10b). These observations confirmed the distinct groupings observed in the MDS embedding of Frobenius distances (Figure 2b).

## DISCUSSION

Data normalization and inference of taxon-taxon associations from microbial genomic survey count data are two of the most basic statistical analysis tasks in modern microbiome research. To help the practitioner of microbial data analysis make informed choices about the different available normalization and association inference schemes, we have taken a closer look at the impact of data normalization on association estimation and several downstream exploratory data analysis tasks. Rather than asking what is the *best* method available for different analysis steps, we have leveraged the large available sample size in the American Gut Project (AGP) data set and assessed the *consistency* of two ubiquitous linear association estimators for microbiome data, correlation and proportionality, under a wide range of realistic sample size scenarios, data normalization schemes, and downstream data analysis tasks.

Our analysis revealed several important observations that have direct implications for best practice in microbiome data analysis workflows. Firstly, we have confirmed that correlation and proportionality estimates are inconsistent in the low sample regime *n* < *p* when compared to large sample counterparts, both in terms of general large-scale association patterns (Figure 2) and downstream network summary statistics, including assortativity and modularity (Figure 7). While this phenomenon has been long appreciated in the statistical literature, we have established that shrinkage estimation, a popular statistical regularization scheme used in finance (28) and genomics research (31), can also improve association estimation for microbiome data, independent of the employed normalization method. Leveraging the close mathematical relationship between variance-covariance estimation and the concept of proportionality, we have also introduced a novel shrinkage proportionality estimator, rhoshrink, that is easy to compute and may prove useful in other scientific areas where compositional data are available.

On the AGP data, we have been able to categorize ten data normalization/association estimation workflows into five coherent groups that show strong agreement across all sample size scenarios (Figure 2b). Most prominently, we have found that variance-reducing/stabilizing transformations lead to a high agreement of correlation or proportionality estimates. This was also confirmed in the downstream microbial relevance network comparison where clr-based correlation networks and proportionality association networks showed high agreement among the inferred edge sets (Figure 8). This implies that, in the presence of large samples sizes and large number of OTUs, differences between correlation and proportionality estimates are less pronounced than previously expected. An important observation on the AGP dataset was that the empirical distributions of association estimates were universally right-skewed even in the randomly shuffled data scenario. This implies that no matter the data normalization/association inference workflow, one will observe a higher prevalence of positive associations. This phenomenon has been previously described in the context of microbial association inference across many different microbial habitats (12, 45). While it is tempting to interpret these results as ecological features of the underlying microbial community in terms of higher prevalence of commensual rather than competitive microbial interactions, the positive skewness may also be due to technical limitations in the data generation process and statistical shortcomings. More specifically, truncation to zero effects for low sequencing read counts likely obstruct unbiased estimation of negative correlations, and in turn, proportionality. A possible remedy for this data-induced artifact is the application of more advanced semi-parametric correlation estimators that infer latent correlations under data truncation assumptions (21, 46). A detailed investigation of semi-parametric and other estimators may provide a promising avenue for future research.

Despite the universal presence of positive skewness in association estimates for the AGP data, we have observed that variance-reducing/stabilizing transformations could reduce positive skewness in shrunk association estimates (Figure 3). Moreover, our results on microbial association network construction and clustering as typical downstream exploratory data analysis examples also revealed that variance-reducing/stabilizing approaches provided the most consistent estimation in terms of taxonomic and structural coherence, as measured by taxonomic cluster purity in spectral and hierarchical clustering (Figures 4 and 5) and network assortativity (Figures 6 and 7). Taken together, we can recommend any variance-reducing/stabilizing transformations followed by shrinkage estimation for association inference. However, transformations such as asinh and clr may be preferred since they are faster to compute than vst, while providing similar statistical properties. The resulting shrinkage correlation estimates can then also serve as input for more involved direct microbial network inference workflows that account for transitive correlations, adjust for additional covariates, or model latent effects (12, 37, 47,48).

For relevance network estimation, consensus network construction, as put forward here for the AGP data (Figure 8), is a straightforward strategy to relax the influence of data normalization. For our AGP consensus network, we found that more than half of the top 2000 edges in the tss-, clr-, rhoprop-, and rhoshrink-based relevance networks were in full agreement, connecting a subset of 346 taxa. The inferred AGP consensus network comprised a majority of positive edges and showed high assortativity at the genus level (0.39) and a maximum modularity of 0.8.

Assortativity increased in the consensus network compared to individual relevance networks. Notably, many taxa in the consensus network were frequently identified as key targets for microbiome therapeutics, such as prebiotic treatment and fecal microbiota transplants, including *Akkermansia muciniphila*, *Prevotella copri*, *Ruminococcus bromii*, and *Faecalibacterium prausnitzii* (49, 50).

Our computational data analysis workflow, available on GitHub and as synapse project (see Data Availability), is fully reproducible, provides all novel shrinkage estimators introduced here, and allows easy extension and comparison to additional data normalization, estimation, and downstream analysis tasks. For instance, future work could include the integration of more advanced zero-replacement strategies (51, 52), application of popular data normalization schemes from single-cell data analysis (53) or the application of other correlation (21, 46) or proportionality estimators, including those available in the propr package (23). Here, rather than using universal thresholding for sparsifying associations, more advanced selection strategies that control false discovery rates (as available in the propr package (23)) may improve the consistency of the microbial association inference workflows.

Going forward, we believe that large-scale reproducible computational analysis workflows that focus on samplesize dependent consistency of statistical estimates are of paramount importance for deriving stable testable hypotheses about the complex interplay between host phenotype and the microbiome from large-scale microbial genomic survey data.

## DATA AVAILABILITY

The code and data used is available as a github repository at https://github.com/MichelleBadri/NormCorr_manuscript and Synapse project syn21654780. Data used for this study was accessed from ftp://ftp.microbio.me/AmericanGut/ag-2017-12-04/. The latest complete American Gut Project dataset can be accessed on Qiita using study ID 10317 (24).

## Supplementary Figures

**Figure 9.**
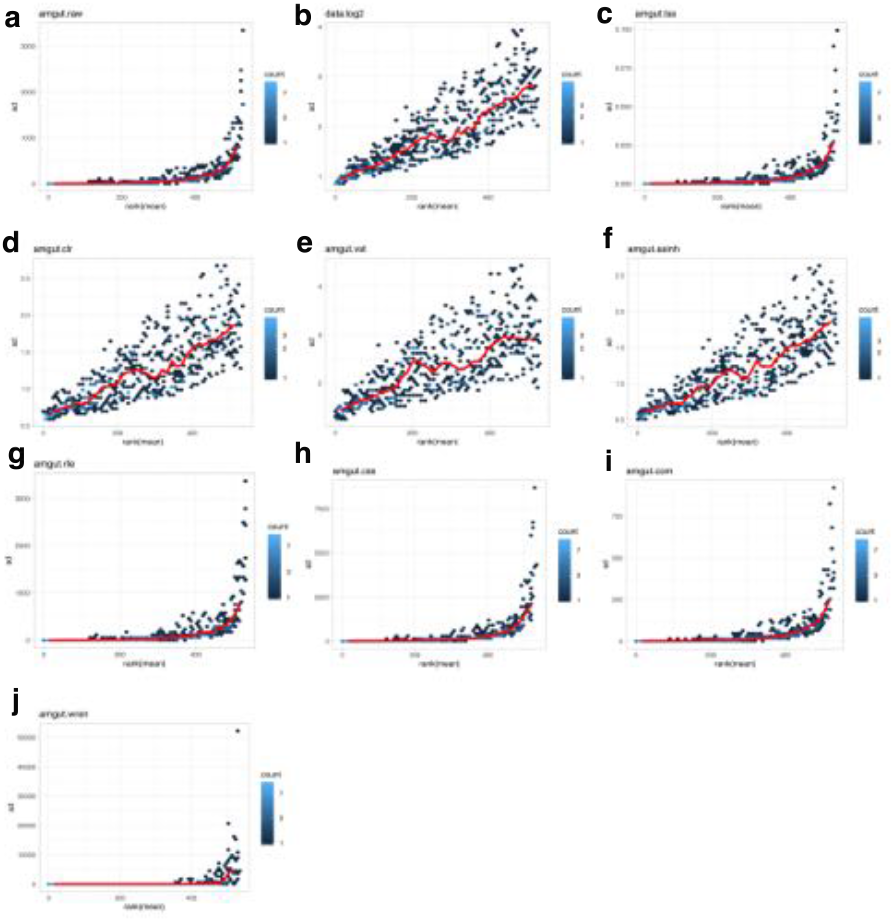
Supplement 1: Standard deviation per-OTU (taken across all 531 OTUs) plotted against the rank of the average count.

**Figure 10.**
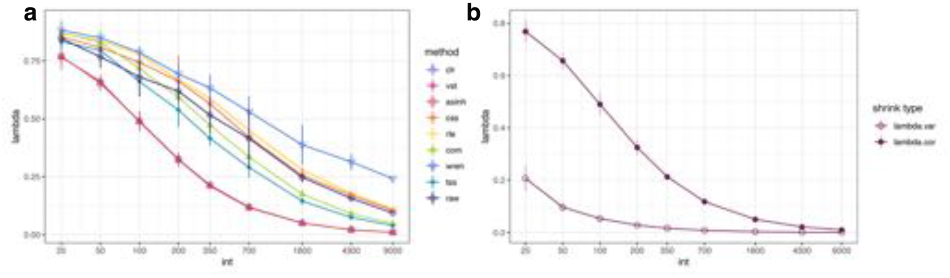
Supplement 2: Lambda values for shrinkage estimation. a) Lambda values selected for shrinkage estimation of correlation 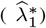. b) Lambda of correlation 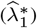 and Lambda of variance shrinkage 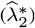 used jointly to estimate shrinkage of covariance for rhoshrink)

**Figure 11.**
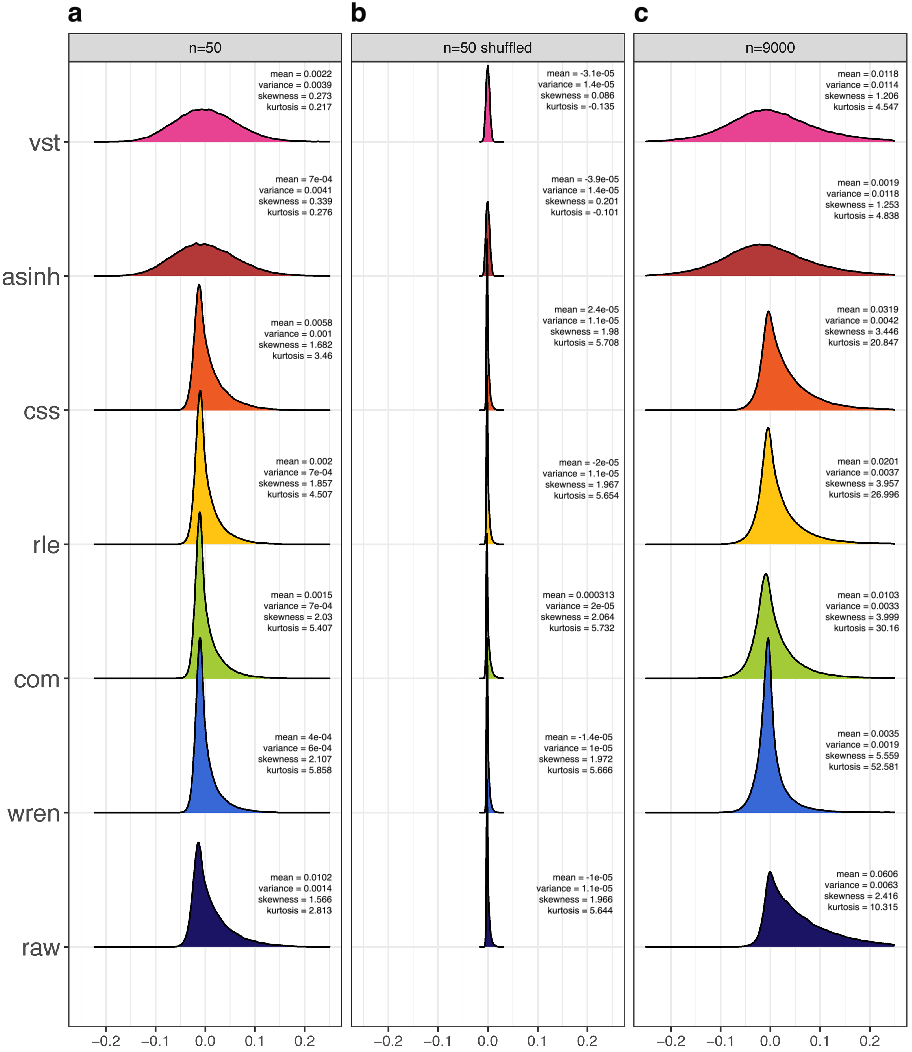
Supplement 3: Frequencies of correlation after transformation. Density plots of correlation frequency of matrices after application of different normalization methods. Mean, variance, skewness, and kurtosis are shown for each distribution.

**Figure 12.**
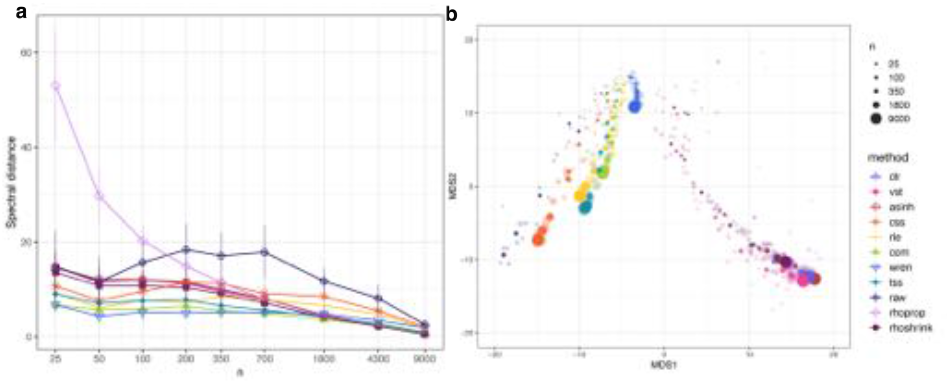
Supplement 4: Spectral distance between estimates of association a) Spectral distance between sub-samples of different sizes. Lines represent mean and error lines represent standard deviation from the mean. Lines represent normalized matrices where correlation/proportionality estimation with shrinkage was performed. b) Multidimensional scaling representation of Spectral distance between correlation structures of varying size estimated from different normalization methods. The most opaque points represent the mean of 5 subsamples of the same size. (color scheme as in A)

**Figure 13.**
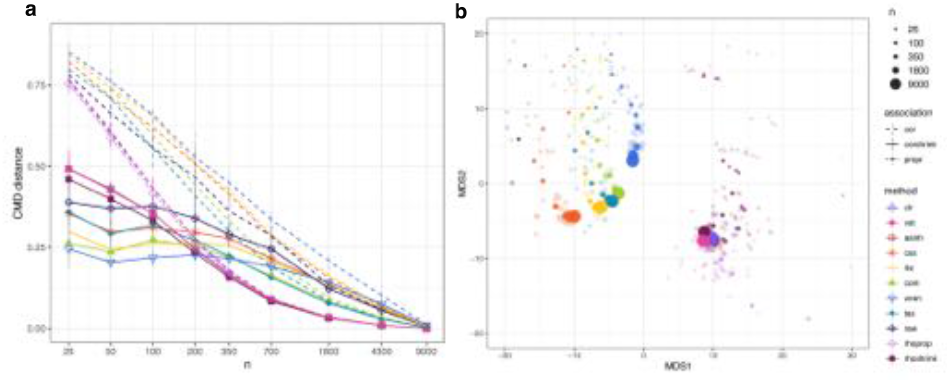
Supplement 5: CMD distance between estimates of association a) CMD distance between sub-samples of different sizes. Lines represent mean and error lines represent standard deviation from the mean. Dashed lines represent normalized matrices after Pearson correlation. The solid lines represent normalized matrices where correlation/proportionality estimation with shrinkage was performed. The dot-dash line represents rho, a proportionality metric. b) Multidimensional scaling representation of CMD distance between correlation structures of varying size estimated from different normalization methods. The most opaque points represent the mean of 5 subsamples of the same size. (color scheme as in A)

**Figure 14.**
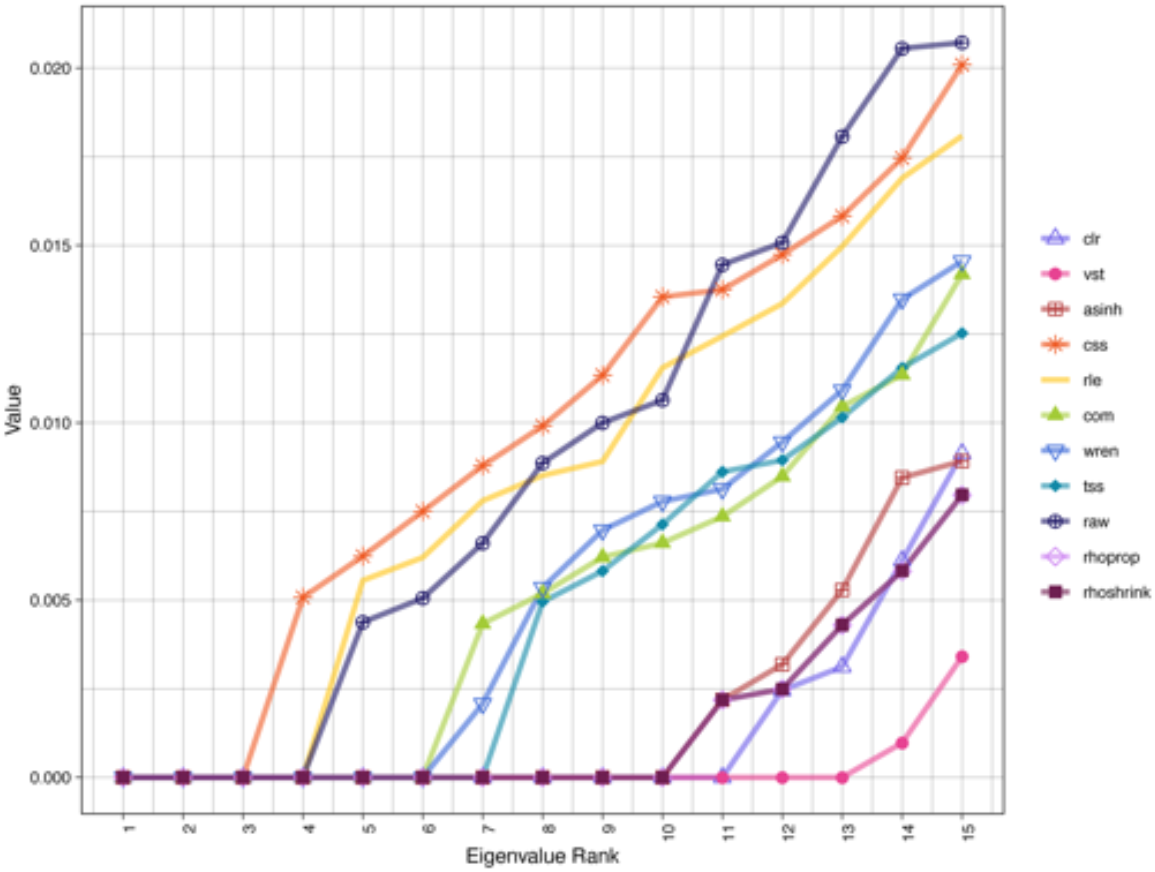
Supplement 6: Numbers of clusters as selected by the number of connected components. Vertical lines represent the first non-zero eigenvalue which is selected as the number of clusters.

**Figure 15.**
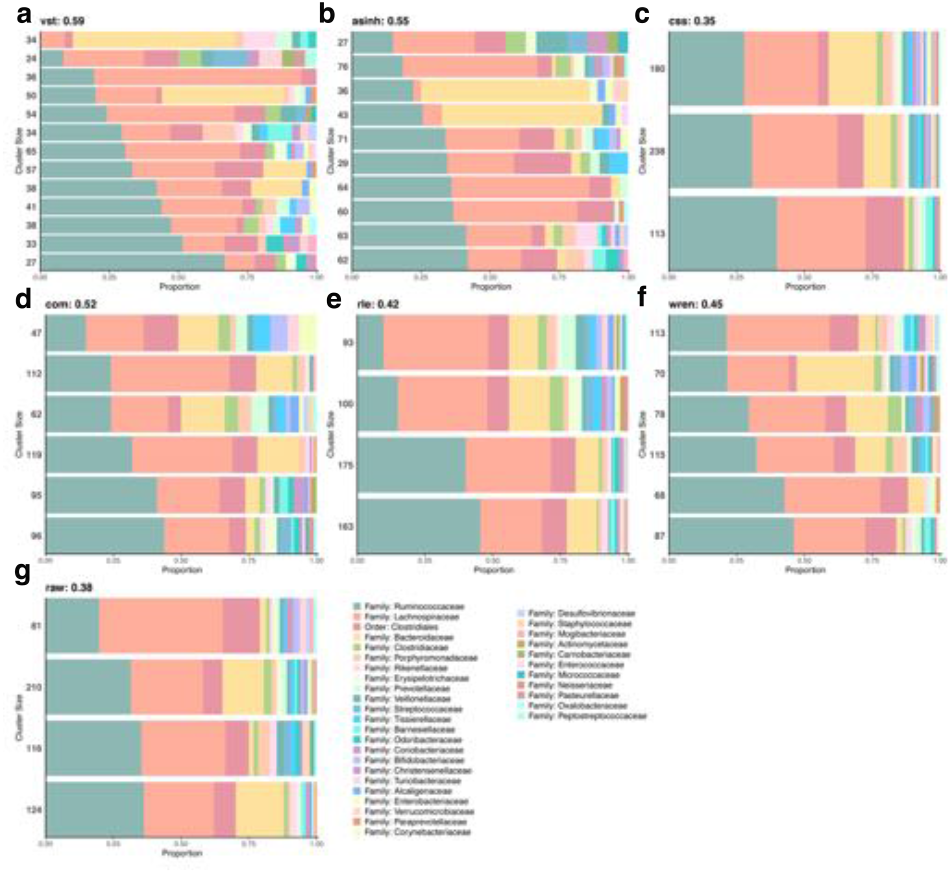
Supplement 7: Horizontal stacked bar plot of OTU groups resulting from spectral clustering. The stacked bar plots represent the composition of OTUs in each cluster at the Family level. Clusters are vertically ordered from highest percentage of the most abundant Family: *Ruminococcaceae*. Horizontally the order represents the highest percentage of Family in each cluster. Numbers on the left axis represent the number of OTUs in each cluster. Values stated next to method name represent cluster purity.

**Figure 16.**
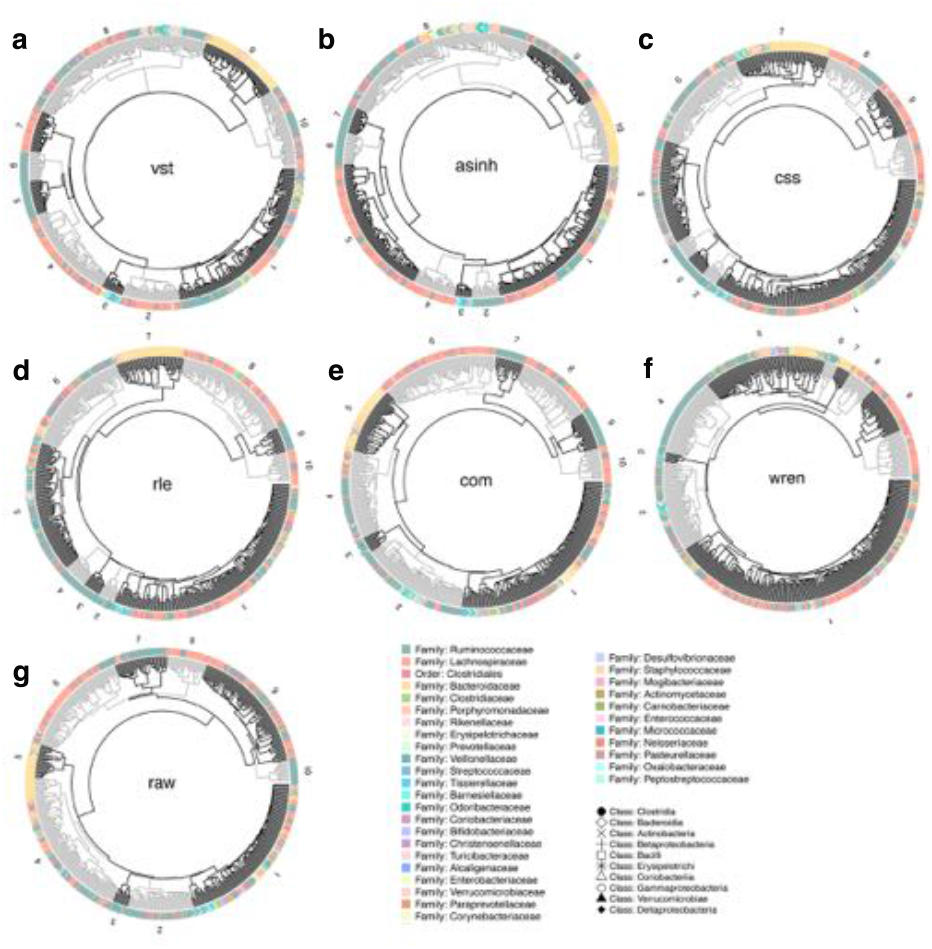
Supplement 8: Circular dendrograms of hierarchical clustering patterns amongst taxa Family and Class. Each point surrounding the circular dendrogram represents one of the 531 OTUs in our data set. The color represents Family annotation and shape represents Class annotation. Each dendrogram has been cut hierarchically into 10 trees (representing the 10 Orders to which each Family maps). The grey and black shading is used to highlight different clusters which are numbered.

**Figure 17.**
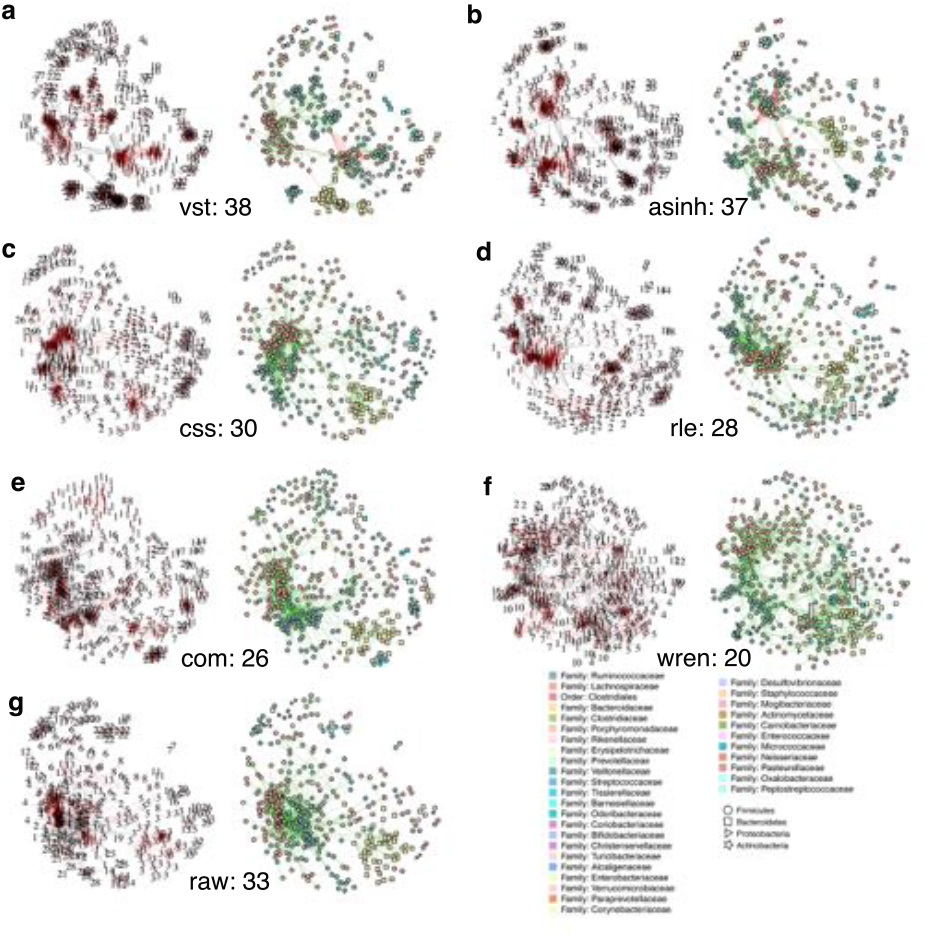
Supplement 9: Relevance network visualization displaying modularity. For networks on the left of each panel every node represents an OTU labelled with module annotation as predicted by the Fast-Greedy modularity algorithm. The networks on the right represent the corresponding phylogenetic annotation of the OTU at the family level. Values stated next to method name represent the number of modules in the network. Layout using the force-directed Fruchterman-Reingold algorithm was conserved for both networks in each panel for comparison

**Figure 18.**
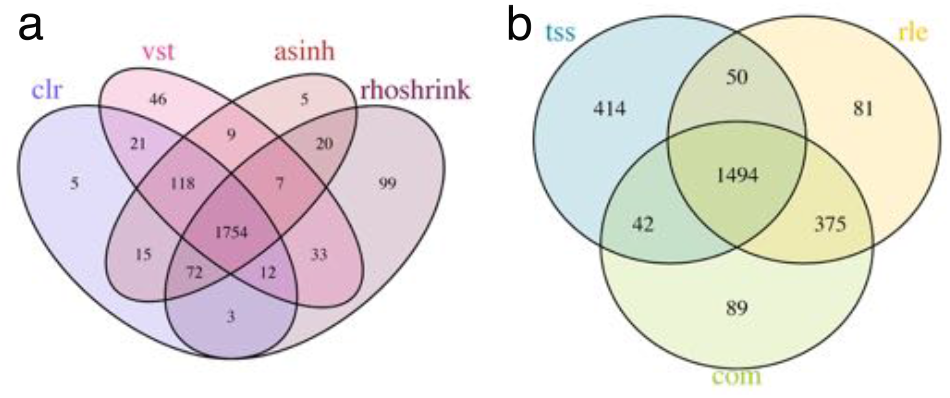
Supplement 10: Edges in common between relevance networks. Venn Diagram of edges in common from the top 2000 edges between a) Log-based normalization methods clr, vst, asinh and rhoshrink b) rle, com, and tss.

## Supplementary Methods

### SHRINKAGE ESTIMATION

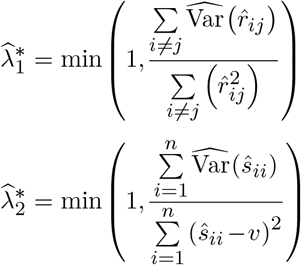

The detailed computation of unbiased estimation of variance 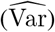 for 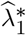 and 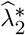 can be found in Schäfer and Strimmer (31, 54).

